# Temporal changes in allele frequency facilitate detection of adaptive variants in winter wheat (*Triticum aestivum* L.) breeding programs

**DOI:** 10.64898/2026.04.30.721918

**Authors:** Natasha H. Johansen, Pernille Sarup, Pernille B. Hansen, Jihad Orabi, Ahmed Jahoor, Guillaume P. Ramstein

**Affiliations:** Center for Quantitative Genomics and Genetics, Aarhus University, 8000, Denmark; Nordic Seed A/S, 8300, Denmark

## Abstract

In quantitative genetics, candidate SNPs are identified through genotype-phenotype associations inferred with genome-wide association studies (GWAS). In this study, we explore an alternative approach to detect genetic variants with non-neutral effects by tracking temporal trends in allele frequency in a winter wheat (*Triticum aestivum* L.) breeding population over an eight-year period, from which signals of selection may be inferred.

Selection signatures were inferred with a generalized linear model, where we modeled trends in allele frequency as a function of time (crossing year). These signatures of selection were used to prioritize variants. Associations between phenotypic performance and individual load of prioritized variants were then investigated. Furthermore, we assessed whether incorporating selection information into a genomic best linear unbiased prediction (GBLUP) model improves model performance in terms of quality of fit and prediction ability.

Our findings indicate that the inferred signals of selection are effective in identifying non-neutral variants. Variants under strong negative selection were associated with a decrease in protein content adjusted for grain yield (p-value < 0.01), while genetic variants that had been under moderate to high levels of positive selection were associated with increased grain yield (p-value < 0.01). However, incorporating selection information did not improve prediction accuracy.

In conclusion, temporal trends in allele frequency can be used to detect non-neutral variants. The proposed approach may hence complement traditional quantitative genetic methods for detecting non-neutral genetic variation. This approach may allow breeders to detect non-neutral variants earlier in the breeding cycle, without resorting to phenotypic data.

## 1. Introduction

Plant breeding aims to accumulate beneficial variants to produce superior breeding material. However, identifying beneficial variants is not a trivial task, as the effect of a variant may depend on genetic context, due to epistasis (Kroymann and Mitchell-Olds 2005; He et al. 2017), or environmental context, due to genotype × environment (G×E) interactions (Lasky et al. 2015; Ferrero-Serrano and Assmann 2019). In quantitative genetics, genotype-phenotype associations are commonly inferred using genome-wide association studies (GWAS). However, methods other than GWAS may be employed to detect non-neutral genetic variation.

These methods include trend tests, which may be used to detect shifts in allele frequency over different time points (Bollback et al. 2008; Feder et al. 2014), through covariance structures (Buffalo and Coop 2020), Bayesian inference (Schraiber et al. 2016), and Hidden Markov models (Mathieson and McVean 2013). These trend tests enable the detection of alleles that undergo a directional change in allele frequency over time, which may be attributed to selection. These methods have the advantage of being computationally less complex than GWAS and do not require phenotypic data. Thus, they may provide an efficient alternative or complement to GWAS. In animal and plant breeding, genomic prediction (GP) models, as first presented by (Meuwissen et al. 2001), are important tools for predicting the genetic merit of candidate lines or breeds (breeding values). GP allows for more efficient detection of superior breeding material compared to phenotypic selection, and as a result, annual genetic gains have increased with the implementation of GP in breeding programs (Tessema et al. 2020).

Multiple studies have reported improved prediction ability of GP models when utilizing selection signatures or fitness-related information to prioritize functionally important variants (Gebreyesus et al. 2020; Long et al. 2022; Wu et al. 2023). For example, (Gebreyesus et al. 2020) presented a novel GP model for the estimation of mortality traits in cattle, where they extended a traditional GBLUP model by accounting for the probability of a calf receiving a double dosage of recessive lethal alleles. Other studies have identified putatively deleterious mutations, based on signatures of selection, which allow for the inference of the burden of deleterious alleles for individual lines or populations. Incorporation of this information in GP modeling has proven advantageous for studies on cassava (Long et al. 2022), potato (Wu et al. 2023), sorghum (Valluru et al. 2019) and maize (Ramstein and Buckler 2022). These studies generally utilized different measures of phylogenetic conservation (PC), which are detected by comparative genetics. These methods are promising to increase genetic gains through genomic prediction, but they may be expensive to implement as they require whole-genome sequencing data (either observed or imputed) and are not applicable to SNP array-based datasets routinely used by breeding companies. Additionally, PC detects long-term evolutionary constraint and is not suited for detection of adaptive variation, i.e., genetic variants associated with increased fitness or improved phenotypic performance under specific environmental conditions or genetic contexts (Mustonen and Lässig 2009; Latrille et al. 2024).

Alternative methods for detection of genetic variants under selection exist that does not require high-resolution sequencing data, including genotype-environment correlations. However, these methods typically require multiple independent populations originating from different environments distributed across an environmental gradient (Coop et al. 2010; De Mita et al. 2013). Breeders generally do not maintain multiple independent populations undergoing selection under distinct environmental conditions within a single program, making this method impractical. Other methods for detecting adaptive variation include F_ST_ tests, which identify genetic differentiation between populations (Akey et al. 2002; Narum and Hess 2011). These tests are similar to trend tests, as both types of methods detect adaptive genetic variation through comparisons among groups of genotypes, but they differ in the patterns of selection they can detect. While F_st_-tests detect differences among discrete populations, trend tests detect continuous variation over time. Previous publications suggest that knowledge about selection signatures may improve GP and the detection of impactful variants (Gebreyesus et al. 2020; Long et al. 2022; Wu et al. 2023). In a breeding context, selection signatures must be inferred in consideration of the constraint of the underlying genomic data while at the same time leveraging the strengths of breeding data, particularly its longitudinal structure, suitable for the inference of temporal trends in allele frequency, thus potentially facilitating the detection of beneficial genetic variants. In this study, we inferred selection signatures based on temporal trends in allele frequency in a breeding program in winter wheat (*Triticum aestivum* L.) over eight years. We then at validating these signatures for their ability to capture non-neutral variants.

## 2. Methods & Materials

### 2.1 Genotyping and imputation

DNA was extracted using the CTAB method (Rogers and Bendich 1985) and genotyped using a 25K Illumina Infinium iSelect HD Custom Genotyping BeadChip (SGS-TraitGenetics, Gatersleben, Germany). Multiallelic sites and sites with minor allele frequency (MAF) below 0.0001 were excluded. Missing genotypes were imputed using BEAGLE v. 5.4 (Browning et al. 2018), with a sliding window of 40 cM and an overlap between adjacent sliding windows of 20 cM. After filtering, 10,230 SNPs remained. The germplasm used in the study was from the Danish breeding company Nordic Seed A/S.

### 2.2 Phenotypic analysis

Different phenotypic datasets were used for GWAS and GP. The GWAS included 959 lines phenotyped between 2015 and 2020, while in the GP analysis we used a temporally separated dataset of 1,225 lines with phenotypic records from 2021 to 2023 **(Figure 1**). Only sixth-generation (F_6_) winter wheat lines were included in both analyses.

**Figure 1:**
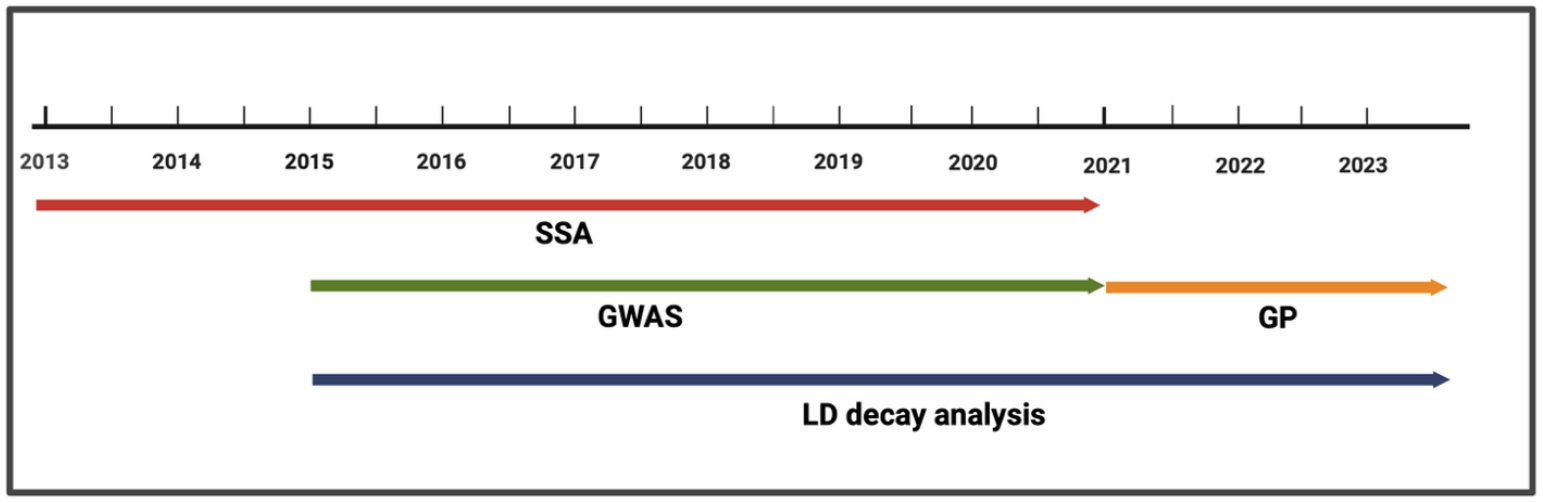
Timeline of crossing, phenotyping years used for downstream analyses. In the selection signature analysis (SSA), signatures were inferred from F_5_ genotypes derived from crosses made between 2013 and 2020. Phenotypic data from F_6_ lines were used for GWAS and genomic prediction (GP) models. Specifically, lines phenotyped between 2015 and 2020 were used in GWAS, whereas lines phenotyped between 2021 and 2023 were used for validation of GP models. All lines included in the GWAS and GP analyses were used for inference of linkage disequilibrium (LD) decay.

Four agronomic traits were included in the study: grain yield, heading date, plant height, and protein content. Grain yield was measured as kg per plot at a moisture level of 15% (plot size: 8.25 *m*^*2*^). Heading date was recorded as the number of days after the 1st of May when spike emergence was observed. It is not optimal to use raw protein content measurements, as yield and protein content are negatively correlated due to physiological constraints in resource allocation. This relationship is observed in multiple cereals, including wheat (Simmonds 1995). Failure to account for this relationship can lead to spurious associations for instance variants associated with high yield may appear to be associated with reduced protein content.

To account for this relationship, grain protein deviation (GPD) was used instead of protein content. This trait was estimated by regressing yield on protein content, where GDP represents the deviation from the yield-protein relationship (Mosleth et al. 2020).

Adjusted entry means were calculated independently for each location-year combination. Linear mixed models (LMMs) were fitted in which entry-level phenotypic records were regressed on genotypes (fixed effects). To adjust for spatial effects at field level, we used a first-order autoregressive correlation structure on field column and field row or alternatively – if this spatial model did not converge – a moving-average structure. Model residuals were assumed to be independent and identically distributed (*i*.*i*.*d*.), following a normal distribution. The adjusted entry means were used as the response variable in downstream statistical models. The adjusted means, *y*_*ijlk*_, correspond to a given genotype/line *i*, country *j*, year *k* and location *l* combination (**Table 1, Figure S1A**).

**Table S1:**
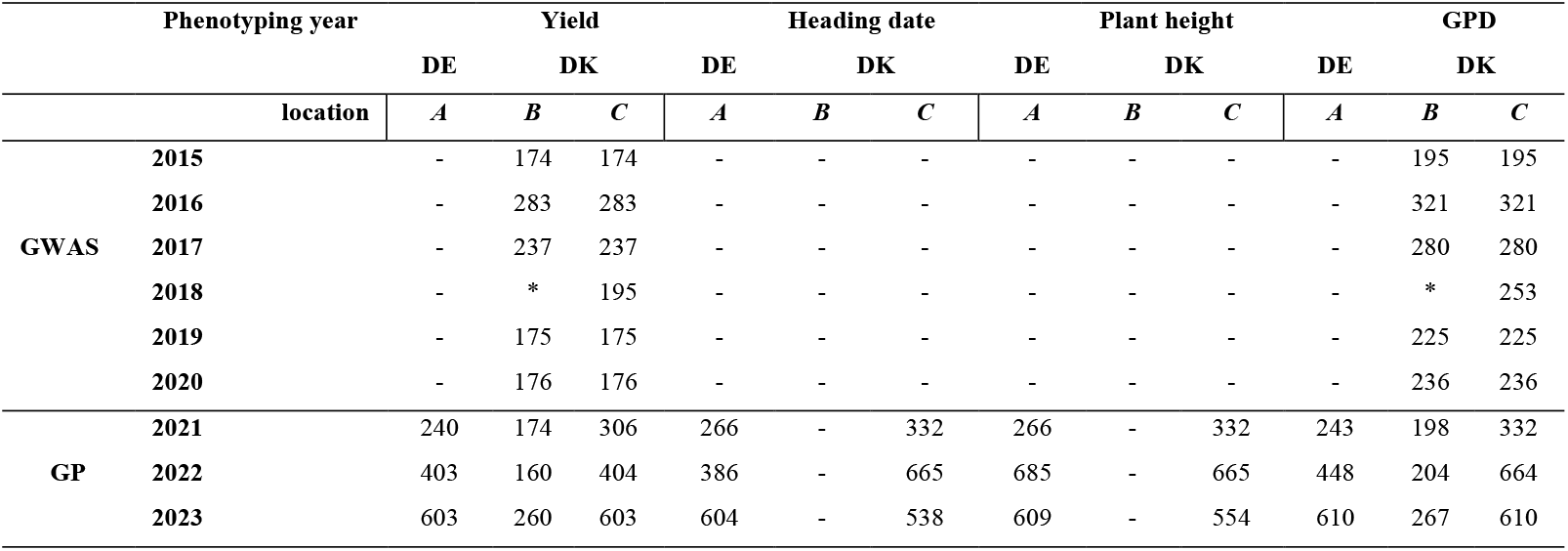
Number of F_6_ winter wheat lines evaluated for each trait across country-location combinations. DK and DE denote Denmark and Germany, respectively. Field trials were conducted at two locations in Denmark: Dyngby (55°57′19.5″N, 10°14′26.0″E), and Holeby (54°42′41.6″N, 11°27′59.4″E), and one location in Germany: Nienstädt (52°17′36.6″N, 9°08′53.6″E). Location codes correspond to A (Nienstädt), B (Dyngby), and C (Holeby). All traits were evaluated at the German site (A), while grain protein deviation (GPD) and grain yield were assessed at both Danish locations (B and C). Heading date and plant height were measured at a single Danish location (C). * Phenotypes not available for location B in year 2018.

| Phenotyping year |  | Yield |  |  | Heading date |  |  | Plant height |  |  | GPD |  |  |
| --- | --- | --- | --- | --- | --- | --- | --- | --- | --- | --- | --- | --- | --- |
|  |  | DE | DK |  | DE | DK |  | DE | DK |  | DE | DK |  |
|  | location | A | B | C | A | B | C | A | B | C | A | B | C |
| GWAS | 2015 | - | 174 | 174 | - | - | - | - | - | - | - | 195 | 195 |
|  | 2016 | - | 283 | 283 | - | - | - | - | - | - | - | 321 | 321 |
|  | 2017 | - | 237 | 237 | - | - | - | - | - | - | - | 280 | 280 |
|  | 2018 | - | * | 195 | - | - | - | - | - | - | - | * | 253 |
|  | 2019 | - | 175 | 175 | - | - | - | - | - | - | - | 225 | 225 |
|  | 2020 | - | 176 | 176 | - | - | - | - | - | - | - | 236 | 236 |
| GP | 2021 | 240 | 174 | 306 | 266 | - | 332 | 266 | - | 332 | 243 | 198 | 332 |
|  | 2022 | 403 | 160 | 404 | 386 | - | 665 | 685 | - | 665 | 448 | 204 | 664 |
|  | 2023 | 603 | 260 | 603 | 604 | - | 538 | 609 | - | 554 | 610 | 267 | 610 |

### 2.3 Statistical models

#### 2.3.1 Selection signature analysis (SSA): Logistic regression to infer signatures of selection

We used a generalized linear model (GLM), with logit link function, to infer signatures of selection, by modelling the trend in allele frequency over time. For inference, we used genotypic information from 18,435 fifth-generation (F_5_) winter wheat lines derived from crosses made between 2013 and 2020. To avoid circularity (i.e., overlap between the data used in trend tests and GP/GWAS), we avoided including lines that had been phenotyped at F_6_ after 2020, as these were used in downstream genomic prediction models for validation. Only F_5_ lines intended for the Danish market were included in the analysis to avoid issues that may arise due to potentially opposing directions of selection in the different environments, i.e., Germany (DE) and Denmark (DK).

Signatures of selection were inferred with the **M0** model:

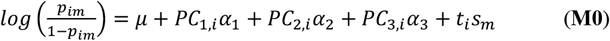

where *p*_*im*_ is allele dosage of the minor allele at locus *m* and line *i* (0 if homozygous for the major allele, 1 if heterozygous, or 2 if homozygous for the minor allele), and *t* refers to crossing year of line *i*. The first three principal components (PCs) from a PCA on the genotype matrix **X** were included to account for individual-specific population structure and to reduce spurious associations resulting from certain genetic clusters being overrepresented in specific crossing years.

The relative advantage or disadvantage of the minor allele compared to the major allele at site *m* was given by the regression coefficient *s*_*m*_. These coefficients were standardized to Z-scores by dividing by the standard error: 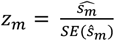. Z-scores above zero indicate that the minor allele increases in allele frequency over time (putatively under positive selection), while negative Z-scores indicate that minor allele decreases in frequency (putatively under negative selection).

For a given threshold *z*, variants with Z-scores above *z* were classified as putatively beneficial, while those with Z-scores below *-z* were classified as putatively deleterious. The quantiles of the absolute Z-score distribution (30%, 50%, 70%, 90%, 95%, and 99%) were used as thresholds classify genomic sites in selection categories for downstream analyses.

#### 2.3.2 Genomic Prediction models

In GP analyses, linear mixed models were used for: (i) **Mean partition:** Investigate the directional effect of genetic variants prioritized by Z-scores. (ii) **Variance partition:** Investigate differential distributions for putatively neutral and putatively non-neutral genetic variants.

##### 2.3.2.1 Genomic relationship matrix

The genomic relationship matrix (GRM) for the additive genetic effect is constructed following the approach described in (VanRaden 2008).

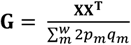

where **X** is the genotype matrix centered to locus means of zero, and ∑_*m*_ 2*p*_*m*_*q*_*m*_, is the scaling factor, with *p*_*m*_ and *q*_*m*_ denoting the frequencies of the minor and major alleles at the *m*^*th*^ locus, respectively.

##### 2.3.2.2 Standard genomic prediction model

The **M1** model is a standard genomic best linear unbiased prediction (GBLUP) model (Habier et al. 2013) that does not incorporate signatures of selection. This model serves as the baseline against which extended models are compared.

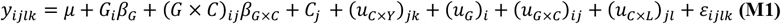

Here *y*_*ijlk*_ is the adjusted entry-mean for line *i*, in country *j*, location *l* and year *k*. The grand mean is denoted by *μ*; the individual genome-wide load of minor alleles is given as *G*_*i*_ = ∑_*m*_ *x*_*im*_, where *x*_*im*_ is the count of minor alleles for line *i* at variant *m*; *β*_*G*_ is its fixed effect, to account for any potential effect of being enriched for minor alleles, while the fixed effect *β*_*G*×*C*_ accounts for country-specific effect of being enriched for minor alleles. The random additive genetic effects for line *i* is given by (*u*_*G*_)_*i*_, where 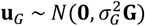, while the G×E interaction term between the additive genetic effect and country is denoted by (*u*_*G*×*C*_)_*ij*_, where 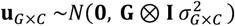. Environmental effects are accounted for by the fixed effect of country, *C*_*j*_, the random effect of year nested in country denoted by 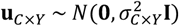, and the random effect of location nested in country given by 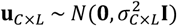. Plant height and Heading date were only scored in a single location in each country, and hence the (*u*_*C*×*L*_)_*jl*_ term was not included when modelling these traits. Lastly, the error term *ε*_*ijlk*_ was modelled as 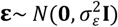, with identity matrix **I**.

Narrow-sense heritability (*h*^2^) measures the proportion of the phenotypic variance explained by additive genetic effects, and was calculated based on variance components from the **M1** model:

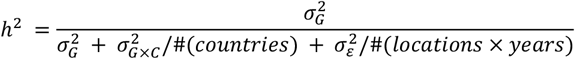

##### 2.3.2.3 Mean partition: genome-wide effects of prioritized variants

The individual loads of putatively deleterious (*D*_*i*_) and putatively beneficial (*B*_*i*_) alleles are determined for all F_6_ lines in the validation set at different thresholds of *z* and −*z*.

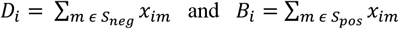

Here *S*_*neg*_ = {*m* ∣ *z*_*m*_ < −*z*} refers to the set of sites where the minor allele is associated with a Z-score *z*_*m*_ below the lower threshold, −*z*, while *S*_*pos*_ = {*m* ∣ *z*_*m*_ > *z*} refer the set of sites where the minor allele is associated with *z*_*m*_ above an upper threshold *z*.

To infer the directional effect of the prioritized variants under negative or positive selection, we fitted model **M1A**, which included the additional covariates *B*_*i*_ and *D*_*i*_ and their fixed effects *β*_*B*_ and *β*_*D*_.

We refer to these fixed effects as the *marginal* effects of prioritized variants, i.e., their average effect across environments (DK and DE).

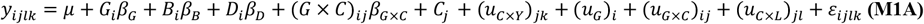

##### 2.3.2.4 Mean partition: genome-wide effects of prioritized variants, including environment-specific effects

To investigate whether the prioritized variants have environment-specific effects we extended **M1A** by including G×E terms: (*B* × *C*)_*ij*_ and (*D* × *C*)_*ij*_ with corresponding fixed effects *β*_*B*×*C*_ and *β*_*D*×*C*_ (Model **M1B**). In the **M1B** model the *β*_*B*×*C*_ and *β*_*D*×*C*_ capture deviations in these effects when comparing Danish environments to the German environments. The effects are referred to as *conditional* effects.

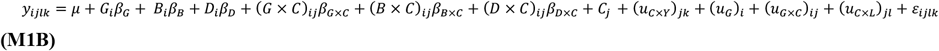

Fixed effects in the **M1A** and **M1B** models were tested using Wald tests (analytical) and permutations tests (empirical) to assess the significance of inferred directional effects of the prioritized variants.

A total of *n* = **250** permutations was used to estimate empirical p-values according to *P* = (1 + *r*)/(1 + *n*), where *n* is the total number of permutations, and *r* the number of permutations fulfilling the significance criterion (i.e., permuted value being equal or more extreme than the observed value, according to the alternate hypothesis) (North et al. 2002). The p-values from the analytical (Wald) test and empirical (permutation) tests are denoted by p_Analytical_ and p_Empirical_, respectively.

##### 2.3.2.5 Variance partition: selection-informed random-effect model

In the selection-informed random-effect GBLUP model **M2**, we investigated whether variants associated with extreme Z-scores, i.e., putatively non-neutral variants in *S*_*neg*_ ∪ *S*_*pos*_, belong to a different effect distribution than the other (putatively neutral) variants.

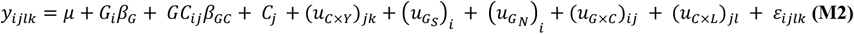

Here we partition the **G** kinship matrix into two kinship matrices; (i) a kinship matrix **G**_*S*_ that describes the relationship among lines based on putatively non-neutral variants, and (ii) **G**_*N*_ based on putatively neutral variants.

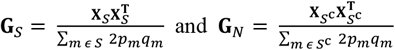

Here the sets of variants are *S* = {*m* ∣ *z*_*m*_ < −*z or z*_*m*_ > *z*} and its complement 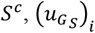 is the additive genetic effect of putatively non-neutral variants, 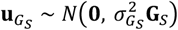, while 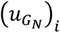 is the additive genetic effect of putatively neutral variants, such that 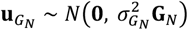.

##### 2.3.2.6 Variance partition: selection-informed random-effect model, including environment-specific effects

The **M2** model was further extended to account for environment-specific variant effects (model **M3**). This extension allowed investigation of whether prioritized variants exhibit different effect distributions between the trial sites in Denmark and those in Germany.

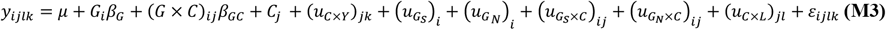

G×E interactions for putatively non-neutral variants are denoted by 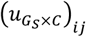, where 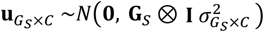, while those for putatively neutral variants are denoted by 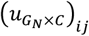, where 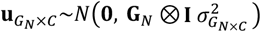.

To test the significance of random effects in variance partition models **M2** and **M3**, we used likelihood-ratio tests (LRTs). Log-likelihood ratios (*LLR*) were computed based on restricted maximum likelihood (REML):

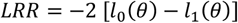

where *l*_1_(*θ*) is the log-likelihood of the complex model, and *l*_0_(*θ*) the log-likelihood of the simpler model, in which the variance of tested random effects is set to zero. The likelihood-ratio test statistic is asymptotically *χ*^*2*^-distributed, with degrees of freedom corresponding to the number of additional variance components in the complex model relative to the simple model.

Permutation tests were also performed as described in Section (***2.3.2.4***). Here, p-values from the analytical test (LRT) and empirical test (permutation) are denoted by p_Analytical_ and p_Empirical_, respectively.

#### 2.3.3 Model comparison

For cross-validation of the GP models, a 10-fold cross-validation scheme was applied. The 1,225 F_6_ winter wheat lines used for GP were partitioned into ten genetic clusters based on genomic similarity. Clustering was performed using hierarchical clustering with Euclidean distance as similarity metric. These clusters were then used to assign the lines into ten groups. The GP models were trained on all but one genetic cluster, which was left out for validation. For each omitted genetic cluster, prediction accuracy was calculated as the Pearson correlation *Cor*(*ŷ*_*ijlk*_, *y*_*ijlk*_) between the predicted phenotypic performance, *ŷ*_*ijk*_, and the adjusted mean phenotypic performance, *y*_*ijlk*_.

#### 2.3.4 Genome-Wide-Association study (GWAS)

To compare the results of a traditional GWAS modeling approach with those of the logistic regression approach for inference of signatures of selection (Section ***2.3.1***), we performed a GWAS using an LMM. The analysis was conducted on F_6_ lines phenotyped between 2015 and 2020 in Denmark (**Table 1**). While signatures of selection were inferred from F_5_ lines crossed between 2013 and 2020, the GWAS was conducted on a distinct set of lines from a similar time-period. This design reflects a realistic breeding scenario in which a breeder may detect non-neutral variants by leveraging either temporal trends in allele frequency or by performing a GWAS on available phenotypic data.

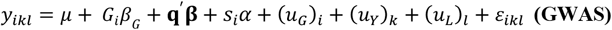

Here *y*_*ikl*_ is the adjusted entry-mean for line *i*, location *l* and year *k*, and *μ* is the overall mean. Population structure was accounted for by including **q**, a vector of the first three principal components for line *i* with corresponding fixed effects **β**. The fixed effect of the tested SNP is represented by *s*_*i*_*α*, where *s*_*i*_ denotes the minor allele dosage for line *i* and *α* is the allele substitution effect.

Random effects included the additive genetic effects 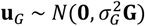, year effects 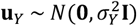, and location effects 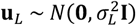 within Denmark. Sites with a MAF below 0.05 were excluded from this analysis.

#### 2.3.5 Linkage disequilibrium analysis

Linkage disequilibrium was calculated as the Pearson correlation (r^2^) as a function of physical distance between loci, with adjustment for population structure. LD decay was thereafter modeled using a generalized linear model (GLM) with a Gamma error distribution and an inverse link function (**Figure S1C**).

#### 2.3.6 Software

For mean partition and variance partition and estimation of likelihoods (REML) we used the R-packages MM4LMM version 3.02 (Laporte et al. 2022) or qgg (Rohde et al. 2020), while adjusted means (BLUEs) was determined with ASREML version (4.2) (Butler et al. 2018).

## 3. Results

### 3.1 Potential signatures of selection are detected by temporal trends in allele frequency

The germplasm used for validation of the GP models was genetically similar and displayed some population structure, as indicated by the first two PCs explaining less than 10 % of the genetic variation (**Figure** 2). Additionally, LD decay analysis showed that the breeding population exhibited relatively rapid LD decay, with LD decreasing below the threshold value, *r*^2^ = 0.20, at approximately 2,500 kbp. Hence, the range of LD blocks in this winter wheat breeding program was relatively short.

**Figure 2:**
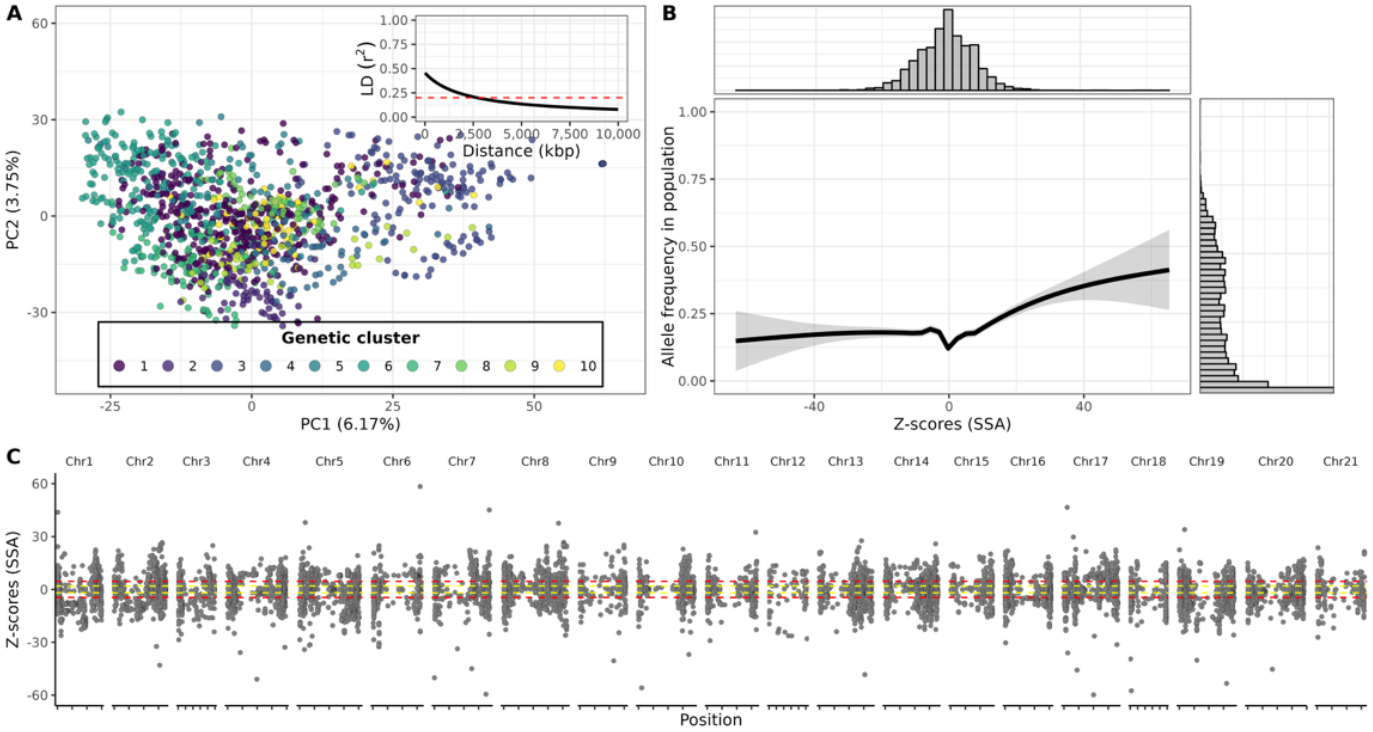
Principal component analysis (PCA) plot of 1,225 F_6_ winter wheat lines used for validation. **A)** Colors indicate assigned genetic cluster. The embedded linkage disequilibrium (LD) decay plot shows the squared Pearson correlation (r^2^) as a function of physical distance within chromosome, specified in kilo-base-pairs (kbp). A threshold at LD, *r*^2^ = 0.20 is indicated. **B**) The upper panel shows the distribution of inferred selection signals (Z-scores) from the selection signature analysis (SSA). The middle panel shows the relationship between Z-scores (SSA) and minor allele frequency, as inferred from a generalized additive model (GAM). The right panel shows the distribution of minor allele frequencies in the breeding program for the lines used for validation. **C**) Genome-wide distribution of standardized signatures of selection (Z-scores; SSA) plotted across genomic positions for allele chromosomes. Each point represents the inferred *z*_*m*_ values for the minor allele at the positions. Positive Z-scores indicate that the minor allele is under positive selection, while negative values indicate the minor allele is under negative selection. The yellow dashed lines indicate significance threshold at the 5% level, not accounting for multiple testing, while the red dashed line indicate the Bonferroni-corrected significance threshold at the 5% level.

We further investigated whether minor allele frequency was associated with inferred selection signatures among the F_6_ winter wheat lines used for validation of GP models. A generalized additive model (Wood 2017) revealed a highly significant association between allele frequency and inferred selection signatures (p < 0.0001, and adjusted R^2^ = 0.055), indicating alleles at higher frequency also tend to coincide with variants inferred to be under positive selection in a temporally distinct dataset.

In the following results, we investigated the prioritization of genetic variants based on inferred signatures of selection. Specifically, we focused on four important agronomic traits with complex genetic architectures that have been under selection in this breeding program. Narrow-sense heritability (*h*^2^) was lowest for production (grain yield, h^2^ = 0.44) and quality (GPD, h^2^ = 0.67), but higher for the physiological traits (plant height, h^2^ = 0.72; and heading date, h^2^ = 0.91) (**Figure S2**). These relatively high heritability estimates suggested that the captured genetic variants account for a substantial proportion of the phenotypic variance, supporting the potential utility of prioritizing these variants based on selection signatures.

From the selection signature analysis, a large proportion of genomic sites are predicted to be under either negative or positive selection (**Figure 2C**). Notably, many selection signals exceeded the significance thresholds (5% significance level). This suggested inflation in the test statistics, which may have led to an overestimation of the number of sites under selection. To avoid, or minimize, potential issues arising from using a hard threshold for classifying sites as being under selection or neutral, we instead applied increasingly stringent thresholds for variant prioritization based on the strength their inferred selection signals. Consequently, as the threshold became more stringent the likelihood of grouping truly neutral and non-neutral sites together decreased, if indeed the inferred signatures of selection corresponded to true signals.

### 3.2 Temporal tendencies in allele frequency detects variants associated with phenotypic performance

In accordance with our hypothesis, we found that variants under negative selection had deleterious effects on at least one agronomic trait (**Figure 3, Table S1**). Specifically, variants predicted to be highly deleterious, those with selection scores exceeding the 99^th^ percentile, were associated with decreased GPD, i.e., lower protein concentration (marginal effect, 99^th^ percentile, p_Analytical_ < 0.01, p_Empirical_ < 0.05, 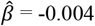, Figure 3A). Furthermore, in line with expectations, variants under apparent positive selection were associated with increased grain yield (70^th^ percentile; p_Analytical_ < 0.01, p_Empirical_ < 0.01, 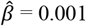, **Figure 3A**).

**Figure 3:**
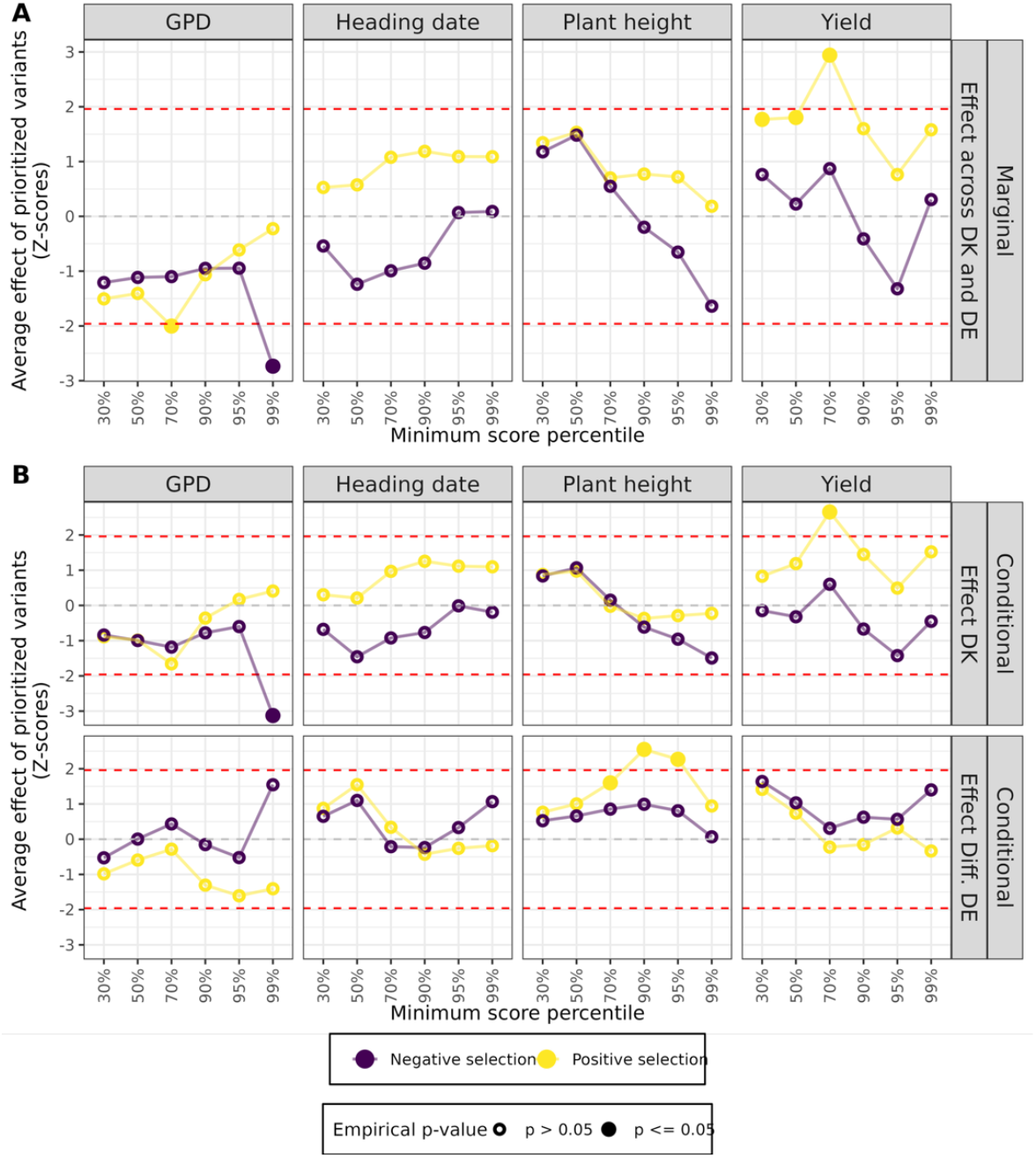
Average effect of prioritized variants across increasingly stringent prioritization of sites (Minimum score percentile) based on inferred signatures of selection. Estimates of the average effect of prioritized variants were standardized by dividing them by their standard errors to obtain Z-scores. Red dashed lines indicate the analytical significance threshold at 5% level (Wald tests), while empirical significance is indicated by filled vs. unfilled dots (permutation tests). A) Results from the M1A model (marginal effects of prioritized variants). B) Results from the M1B model (effect of prioritized variants in DK, and differential effects between Germany and Denmark).

We also investigated whether the prioritized variants exhibited environment-specific effects, i.e. differential effects across environments. Variants that appeared to have non-neutral effects in **Figure 3A** generally showed no significant deviation in effects between Danish and German trial sites, with one exception shown in **Figure 3B**: variants under positive selection were associated with increased plant height in the German trial sites (conditional effect, 90^th^ percentile; p_Analytical_ < 0.01, p_Empirical_ < 0.01, 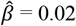, **Figure 3B**).

Overall, the results of mean partition analyses suggest that the effects of prioritized variants are generally consistent across environments for GPD and grain yield and that they may capture biologically meaningful signals. Because signatures of selection were inferred from genetic data of winter wheat lines intended for Danish environments, it might be expected that these signatures are most strongly reflected in the Danish trial sites, this does not appear to be the case.

Stringent prioritization based on inferred signatures of selection, i.e., *negative* Z-scores exceeding the 99^th^ percentile, was associated with a least one trait, whereas prioritization based on the most extreme positive Z-scores (above the 99^th^ percentile) showed no consistent association with any tested trait. This may reflect practical constraints in selection on rare favourable alleles, as in this scenario breeders may be limited in their ability to exert strong selective pressure for these alleles, as strong selection on low-frequency alleles can reduce genetic diversity. Alternatively, many alleles under strong positive selection may already segregate at high allele frequency due to long-term selection in the breeding population. Thus, beneficial variants may seldomly occur at low allele frequencies, potentially when they are recently introduced into the breeding population, either through *de novo* mutations, or via the introduction of foreign breeding material. It is also possible that variants under strong positive selection may be associated with traits not evaluated in this study.

### 3.3 Variant effect distributions differ between putatively neutral and putatively non-neutral variants

We hypothesized that the distribution of variant effects would differ between sites predicted to be neutral and those predicted to be non-neutral based on inferred signatures of selection. To test this, we extended the baseline GBLUP model, which typically includes a single kinship matrix, into a model with two kinship matrices (M2), allowing for separate modelling of random effects of variants classified as neutral and non-neutral. This approach assumed that variants under selection contribute more to phenotypic performance than variants not under selection. M3 further extends this by modelling kinship-specific interactions with environment (country).

The **G**_*N*_ and **G**_*S*_ kernels, representing relationships based on non-prioritized and prioritized variants respectively, showed high correlation among their off-diagonal elements at the 30-70% percentile thresholds, and decreasing to moderate levels for the 90-95% percentiles, and dropping below 0.3 at the 99% percentile threshold (**Figure S1B**). This indicated increasing divergence between the two kernels as more stringent thresholds are applied, suggesting that as the lower threshold for partitioning variants into neutral and non-neutral sites becomes more stringent, increasingly distinct genomic relationship structures are captured.

Results shown in **Figure 4** and **Table S2** indicated that there is no benefit in modeling different variant effect distributions for the physiological traits (Heading date and Plant height). However, for grain yield and GPD, there is strong evidence that variants under selection (GPD: 90^th^ percentile and above) and (grain yield: 95^th^ percentile and above) follow a different effect distribution than putatively neutral variants. For GPD, we additionally observe that the interaction term with environment improved model fit, suggesting that environment-specific interactions differ between putatively neutral and non-neutral variants. Consistent with our expectations, comparison of the variance components showed that variants under selection explained a larger proportion of the phenotypic variance for GPD and yield than putatively neutral variants (**Figure S3**). However, despite convincing improvements in model fit with the extended GBLUP models M2 and M3 (**Figure 4**), no corresponding improvement in genomic prediction accuracy was observed based on leave-one-genetic-cluster-out cross-validation (**Table S3**).

**Figure 4:**
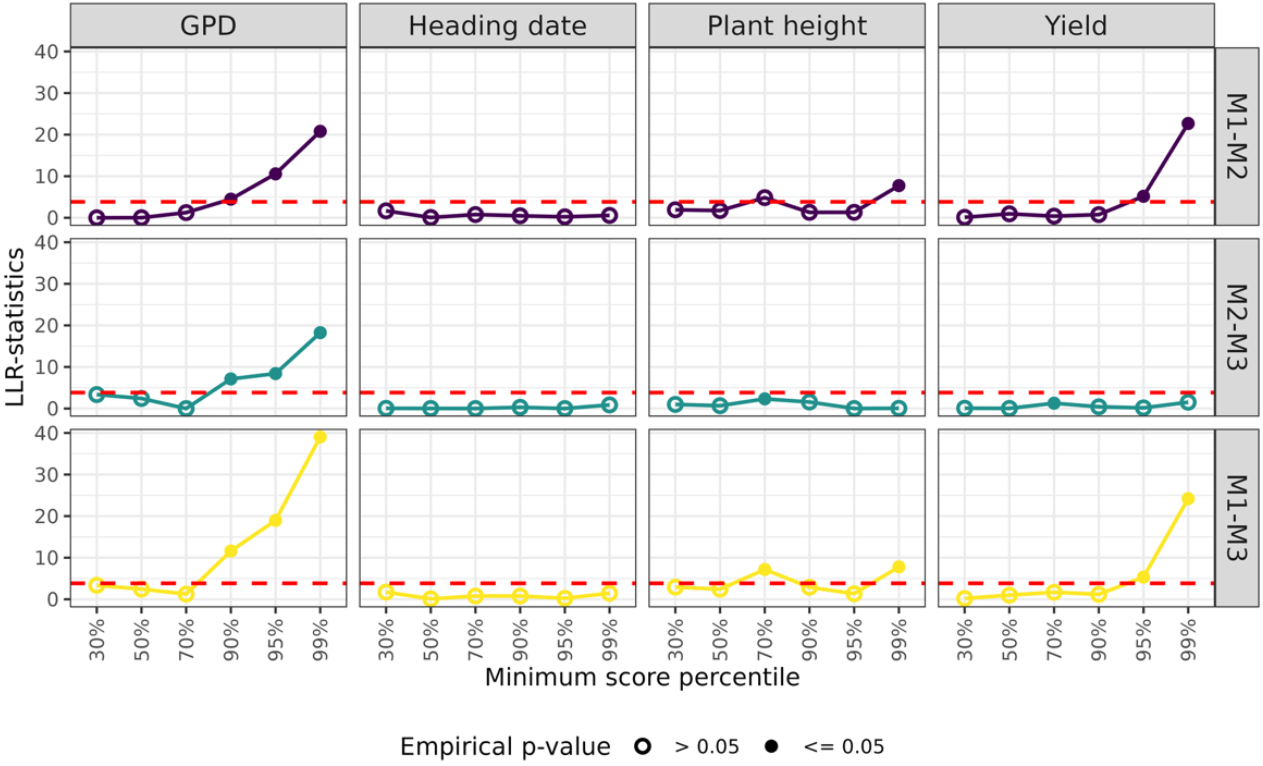
Log-Likelihood ratio (LLR) statistics from nested linear mixed-models show that selection-informed GBLUP model experience improved model fit for Yield and GPD. **M1**: Baseline GBLUP model, **M2**: Selection-informed GBLUP model, and **M3**: Extended selection-informed GBLUP model assuming environment-specific variant effects for the different kinship matrices Red dashed lines indicate analytical significance at the 0.05 level, while filled dots indicate empirical significance as assessed with permutation tests, with n = **250** permutations.

### 3.4 Potentially impactful variants may be identified through GWAS or selection signature analysis

Previous analyses (**Figures 3-4**) indicated that the inferred signatures of selection can be useful for identifying impactful variants for field traits. However, the strength of selection acting on alleles contribution to the tested traits varies, and therefore trait-specific lower thresholds are required when partitioning sites as under selection or not. We also found that many sites appeared to be under directional selection, i.e., p-values exceed the 5% significance threshold (**Figure 2C**). This pattern contrasts with the relatively few significant associations observed in the GWAS results (**Figure 5, Table S4**).

**Figure 5:**
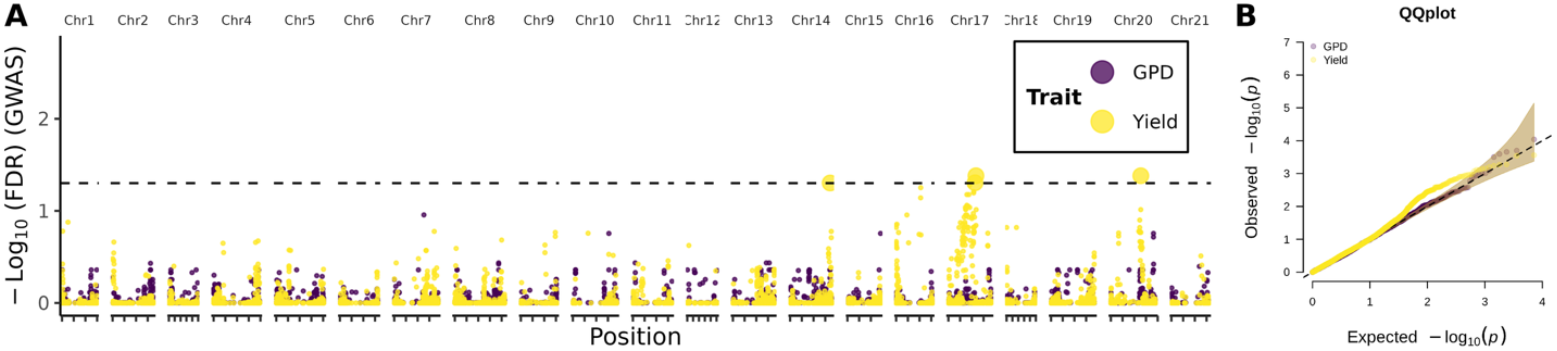
Genome-Wide Association study of important agronomic traits. **A)** Results from Genome-Wide Association studies (GWAS) of grain protein deviation (GPD) and grain yield. To account for multiple testing, p-values were transformed into false discovery rate (FDR)-adjusted values and expressed as −log_10_(FDR). The horizontal line indicate significance at the 5% level. **B**) Quantile-Quantile (QQ) plot shown for the GWAS analyses.

However, signatures of selection may also capture signals linked to traits not investigated in this study, which nevertheless contribute overall agronomic performance. Therefore, the discrepancy between the number of significant associations for agronomic traits (GWAS) and the signatures of selection analysis (SSA) does not necessarily imply inflation in the signatures of selection (**Figure 2C**).

The prioritization approach used in the current study (**Figures 3-4**) did not rely on a single hard significance threshold but instead evaluated a range of thresholds for partitioning of sites into putatively beneficial or deleterious sites (mean partition) or into putatively neutral and non-neutral sites (variance partition). This approach appeared to be more suitable than a single hard threshold, potentially because it reduced the probability of including effectively neutral sites in SNP prioritization as the threshold used grew more stringent.

Finally, we assessed the concordance between signals from SSA and GWAS (**Figure 6**). A consistent positive correlation was observed across thresholds in Z-scores from both analyses, except for the most stringent percentile thresholds (95% and 99% for GPD, and 99% for Yield). This suggests that relying on GWAS alone would not have identified the impactful variants for GPD with negative effects on GPD in both Danish and German trial sites. Highlighting the added value of incorporating signatures of selection for variant identification.

**Figure 6:**
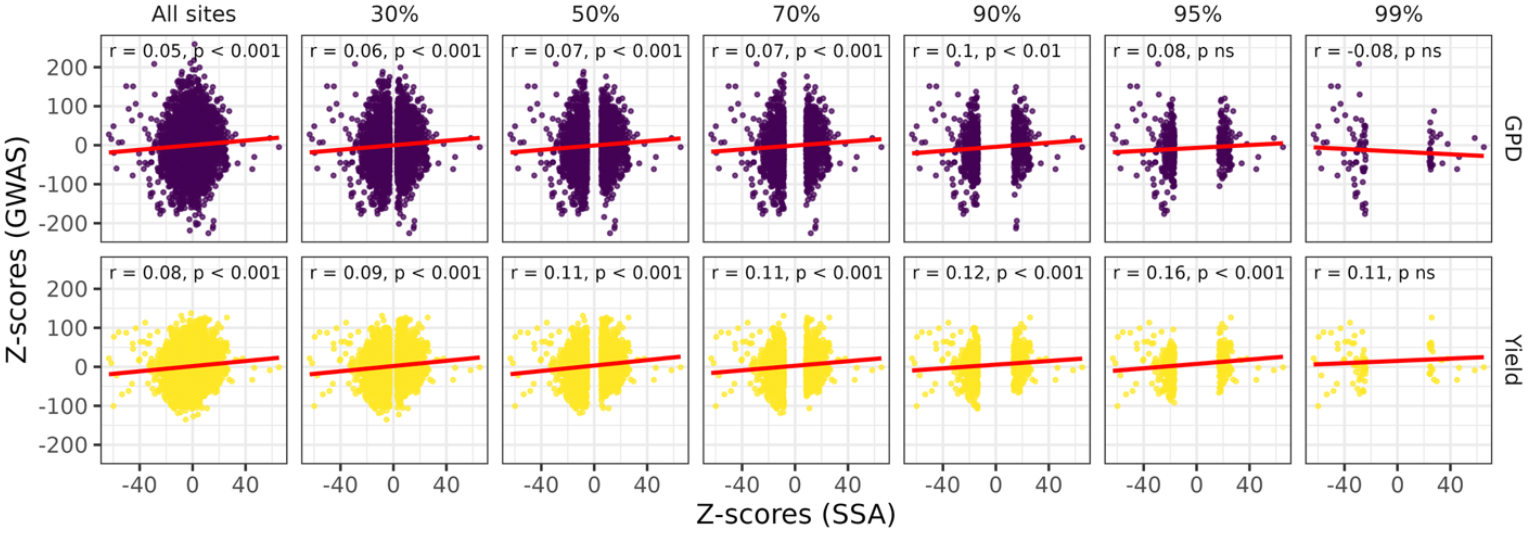
Pearson correlations between inferred signatures of selection and estimated effects from a GWAS. Pearson correlation between standardized signatures of selection (Z-scores; SSA) and standardized effect estimates (Z-scores; GWAS) are shown for increasingly stringent lower thresholds of *z*.

## 4. Discussion

### 4.1 Breeding programs, and the use of trend-tests to detect non-neutral variants

Previous studies have used temporal trends in allele frequency to detect non-neutral variants, tracking shifts in allele frequency over extended time scales or across many generations (Schraiber et al. 2016; Buffalo and Coop 2020; Akbari et al. 2024). However, plant breeding companies often lack historical data spanning many generations. Here we demonstrate that even with a limited number of time-points, trend tests can successfully identify beneficial and deleterious variants impactful for important field traits (**Figures 3-4** and **6**). Efficient identification of variants under selection using the trend-test approach can support breeders in at least two ways: (i) it allows them to verify that the genomic regions under selection are associated with improved phenotypic performance, and (ii) it facilitates detection of favourable genetic variation that may be missed by GWAS. Additionally, unlike GWAS, trend tests are not limited to specific traits and therefore does not disregard loci without direct association to breeder scored traits that may nonetheless be adaptive or favourable.

In this study, we identified multiple significant associations between phenotypic performance and the load of putatively non-neutral variants. Specifically, we showed that variants under strong negative selection were associated with reduced GPD, while those under moderate positive selection were associated with increased grain yield (**Figure 3**). Nevertheless, we did not observe an increase in prediction accuracy relative to a baseline GBLUP model when partitioning the variance of variant effects based on selection signatures (**Table S3**). However, other studies have reported that variance partitioning based on selection signatures – detected by comparative genomics – does translate to increased PA under leave-one-genetic-cluster-out validation schemes (Ramstein and Buckler 2022). Overall, this suggests that prioritization of variants based on selection signatures may offer advantages, but this benefit may depend on the approach used to detect selection signatures, genotyping assay (e.g., SNP array or whole-genome sequencing), validation scheme, and species/populations.

### 4.2 The usefulness of trend tests depends on the genetic architecture?

The traits investigated in this study (heading date, plant height, grain yield and GPD) are all polygenic, meaning many alleles of small effect contribute to these traits. Polygenic selection is often characterised by subtle shifts in allele frequency at many loci, whereas simpler traits are shaped by selective sweeps, i.e., rapid increases in frequency of beneficial alleles until fixation (Stephan 2016; Barghi et al. 2020). This suggests that variants associated with strong selection signals may be associated with traits with simple genetic architectures. In line with this, we observed that variants associated with moderate selection signals increased grain yield (**Figure 3**), a highly polygenic trait (Cao et al. 2020).

Despite moderate to high narrow-sense heritability for the physiological traits (plant height and heading date; **Figure S2**), incorporating selection signals did not improve model performance, i.e., model fit or predictive ability (**Figure 4, Table S3**). In contrast to yield and GPD, which exhibited lower heritability (**Figure S2**), selection-informed models experienced an improvement in model fit (**Figure 4**), but not in predictive ability (**Table S3**). This pattern may reflect differences in the intensity and consistency of breeder-driven selection and highlights that, even when the genetic architecture of a trait aligns with this approach ultimately the detected signals depend on past selective pressures imposed by breeders and natural selection. These selective pressures may arise from both direct selections, e.g., on GPD or yield or indirectly on correlated traits e.g., disease resistance traits. Hence, given that disease resistance is characterised by few high-effect QTLs (Häberle et al. 2009; Miedaner et al. 2020), trend tests may be particularly sensitive to long-term selection on these traits.

The comparatively lower narrow-sense heritability for GPD and grain yield suggests that the genetic architecture of these traits is also largely controlled by non-additive genetic effects, and indeed winter wheat is a crop characterized by extensive epistatic interactions, i.e., gene-by-gene interactions (Raffo et al. 2022). In that respect, it may be advantageous to investigate whether the loci identified as being under directional selection are subject to strong gene-by-gene interactions, to ensure that the identified non-neutral variants are consistently favourable or unfavourable across different genetic backgrounds present in the specific breeding program. If that is not the case, it may in the future be useful to investigate methods for detecting co-selected variants, e.g., genetic background-dependent increases in allele frequencies, as this may be used to detect epistasis. However, due to LD, it would be challenging to distinguish between variants which increase together due to haplotype structure versus those that co-occur due to favourable epistatic interactions. Potentially by focusing on variants located on different chromosomes, it may be possible to detect variants that appear to be jointly favourable by selection (genetic background *sensitive*) and those that are independently favourable (genetic background *insensitive*).

### 4.3 Selection signatures are robust under homogeneous selective environments

Our results indicate that, in this specific breeding program, trend tests are suitable for detecting alleles with environment-independent effects across the tested environments. This is even though the signatures were inferred exclusively based on genetic information from lines intended for the Danish market. This suggests that the variants under selection have similar effects in both production environments, potentially because the environments are so homogeneous that variants conferring an adaptive advantage in Denmark are also favourable in Germany. This highlights the importance of carefully considering the structure of the germplasm used for inference of selection signatures. If there is evidence that the breeding population comprises distinct clusters experiencing differential selection, e.g., as initially hypothesized for Danish and German production environments, selection signatures inferred at the whole-population level may be less robust. Therefore, when applying trend test to infer selection signatures, breeders should remain mindful of past selective environments, as unrecognized genetic or environmental heterogeneity may dilute true signals of selection.

### 4.4 Considerations of how linkage shapes selection patterns

Generally, if the breeding germplasm is characterized by short-range LD, the association between the causal variant (under selection) and the tracked variant may break down over time. Strong LD may hence be favourable for these tests, especially if trends are tracked over many generations, which increases the probability of the causal variant being decoupled from the tracked variant due to recombination events (Buffalo and Coop 2020). On the other hand, LD can introduce interference from non-neutral variants linked to variants under selection. If the variant under positive selection is tightly linked to unfavourable variants, the effectiveness of selection will be diminished. Conversely, if the variant under selection is linked to multiple favourable genetic variants it can enhance selection efficiency (Hill and Robertson 1966). Consequently, beneficial variants in tight LD with unfavourable genetic variation may remain undetected with the current approach (repulsion phase). Meanwhile, strong signals of selection could be a result of tight linkage between multiple variants of similar effects which would increase the selection efficiency (coupling phase). The winter wheat breeding population analysed in this study was genetically homogenous with limited population structure and a moderate rate of LD decay (**Figure 2**), with LD decaying to r^2^ = 0.2 at approx. 2,500 kbp. Other studies report LD decay rates at a physical distance of 11,800 kbp for durum wheat (Roncallo et al. 2021) and 4,980 kbp in winter wheat (Ladejobi et al. 2019). Overall, the size of the linkage blocks and the rate of LD decay should be considered in combination with the number of time points (cross-over opportunities) used for inference of selection signatures. Rapid LD decay over long time frames may reduce robustness. This is unlikely to pose an issue in our study, due to size of the LD blocks and the short time frame (few cross-over opportunities). However, for other crops characterized by rapid LD decay, such as maize (Remington et al. 2001) and perennial ryegrass (Fè et al. 2015), this may be a more important limitation, and higher SNP densities may be required to ensure more robust inference of selection signatures in these populations/species.

### 4.5 Combing selection signature analyses with traditional QTL mapping approaches in breeding programs

The trend tests may, due to practical constraints, be unsuitable for detecting rare beneficial variants, as breeders need to consider genetic diversity and thus cannot exert strong selective force on very rare alleles. Similarly, GWAS often fails to detect true significant associations when the MAF is low, as the sample size has to be quite large for the test to have enough statistical power to detect a significant association (Tam et al. 2019). It is uncertain whether trend tests may perform better in such scenarios where MAF is low, and further investigation would be advantageous to compare trend tests and GWAS for variant discovery when MAF is low. However, as the trend test does not require phenotypic information higher sample sizes may be achieved for this type of test, which may be an advantage when MAF is low, potentially particularly when the deleterious variants is the minor allele.

### 4.6 Conclusions

Trend tests may be used in combination with traditional quantitative genetics approaches, e.g. GWAS, to detect impactful genetic variation. If the breeding goal is to produce material adapted to specific growing conditions, i.e. environmental conditions experienced in Danish and German production environments, trend tests will be more suited than comparative genomic approaches, as trend tests may be used to detect variants with context-dependent effects, i.e., variants adaptive in the intended production environments. Furthermore, trend tests may be a useful tool for breeders to verify that deleterious genetic variation has been removed and given that the method does not require extensive phenotypic data it is a comparatively efficient method for detection of putatively non-neutral variants which may facilitate more rapid improvement of breeding material.

## Acknowledgements

We thank the breeding company (Nordic Seed) for providing access to the plant material and field data used in this study. We also acknowledge the technical staff involved in phenotyping and data management.

## Declarations

### Data and code availability statement

The data supporting the findings of this study is not publicly available due to restrictions imposed by the breeding company in which the data were generated. The analysis code used in the study is publicly available at https://github.com/TashaDear/Signatures_of_selection_winterwheat.git.

### Author contributions

PS, NHJ, and GPR conceptualized the study. NHJ and GPR performed the data analyses. The breeding lines and field data were generated within the breeding company (Nordic Seed A/S), by AJ, JO, PBH and PS. NHJ drafted the manuscript with input from all authors. All authors contributed to interpretation of results and approved the manuscript.

## Supplementary material

**Figure S1:**
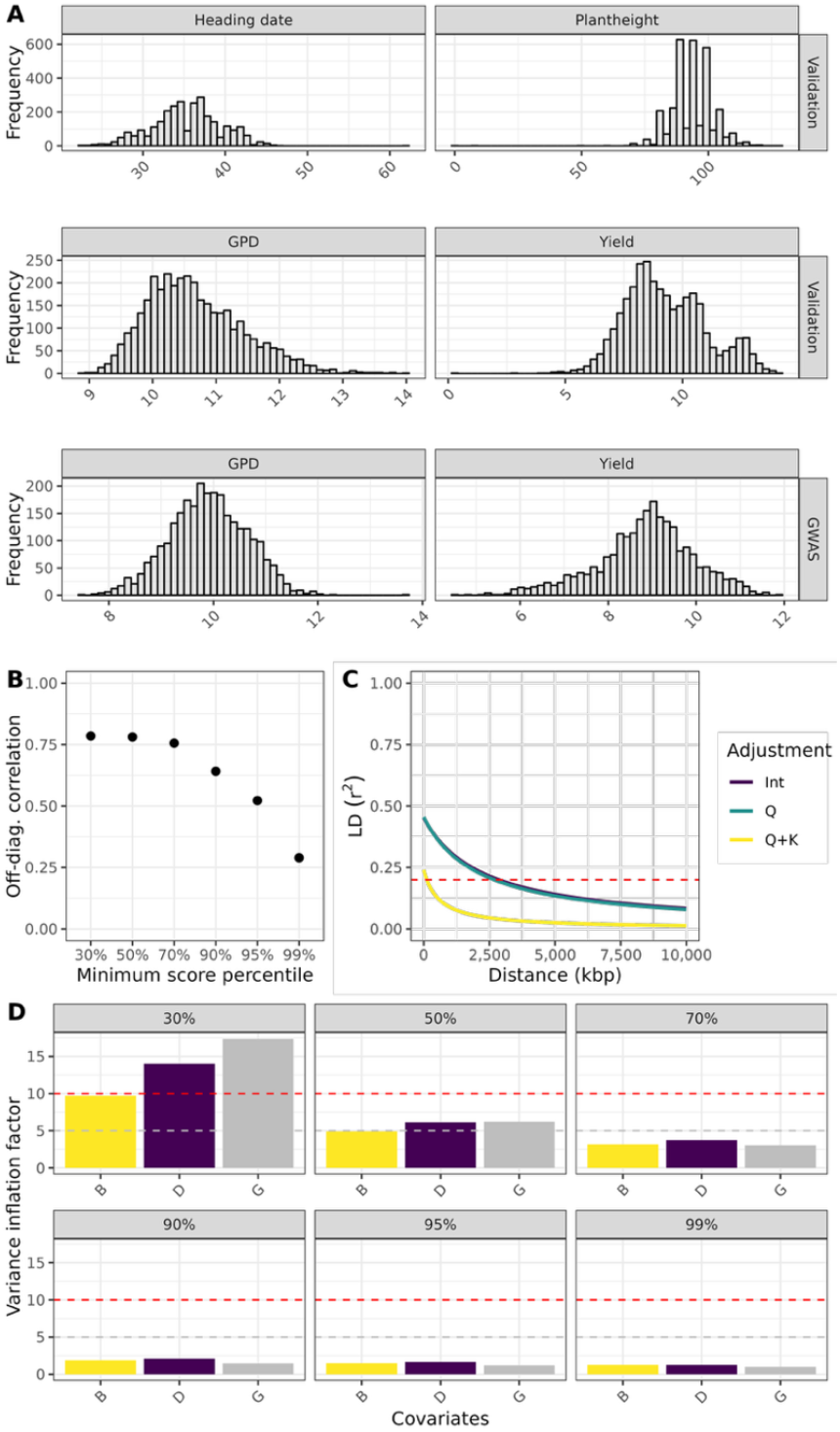
Distribution of phenotypes and linkage disequilibrium. **A)** Distribution of line means, adjusted for spatial effects in the training sets of genomic prediction models (validation) and for the temporally distinct dataset used for GWAS. **B**) The correlation between the off-diagonal elements of two GRMs: **G**_*G*_ and **G**_*L*_ at increasingly stringent prioritization of variants. **C**) Rate of LD decay shown for three different models, (Int) No adjustment for population structure of genomic relationship, (Q) Adjustment for population structure, (Q + K) Adjustment for population structure of genomic relationship. **D**) Variance inflation factors (VIF) used to assess multicollinearity among fixed effects in the mean partition models (M1A). The genome-wide load (G) represents the load of minor alleles, while B and D are the count of putatively beneficial (positive selection) and putatively deleterious (negative selection) minor alleles at a given Z-score threshold for *z*. Horizontal lines indicate common threshold values.: VIF below 5 suggests low multicollinearity, while VIF between 5 and 10 indicates moderate collinearity, and VIF above 10 indicates high collinearity. The level of collinearity observed at the 30% and 50% percentiles may challenge the mean partition models.

**Figure S2:**
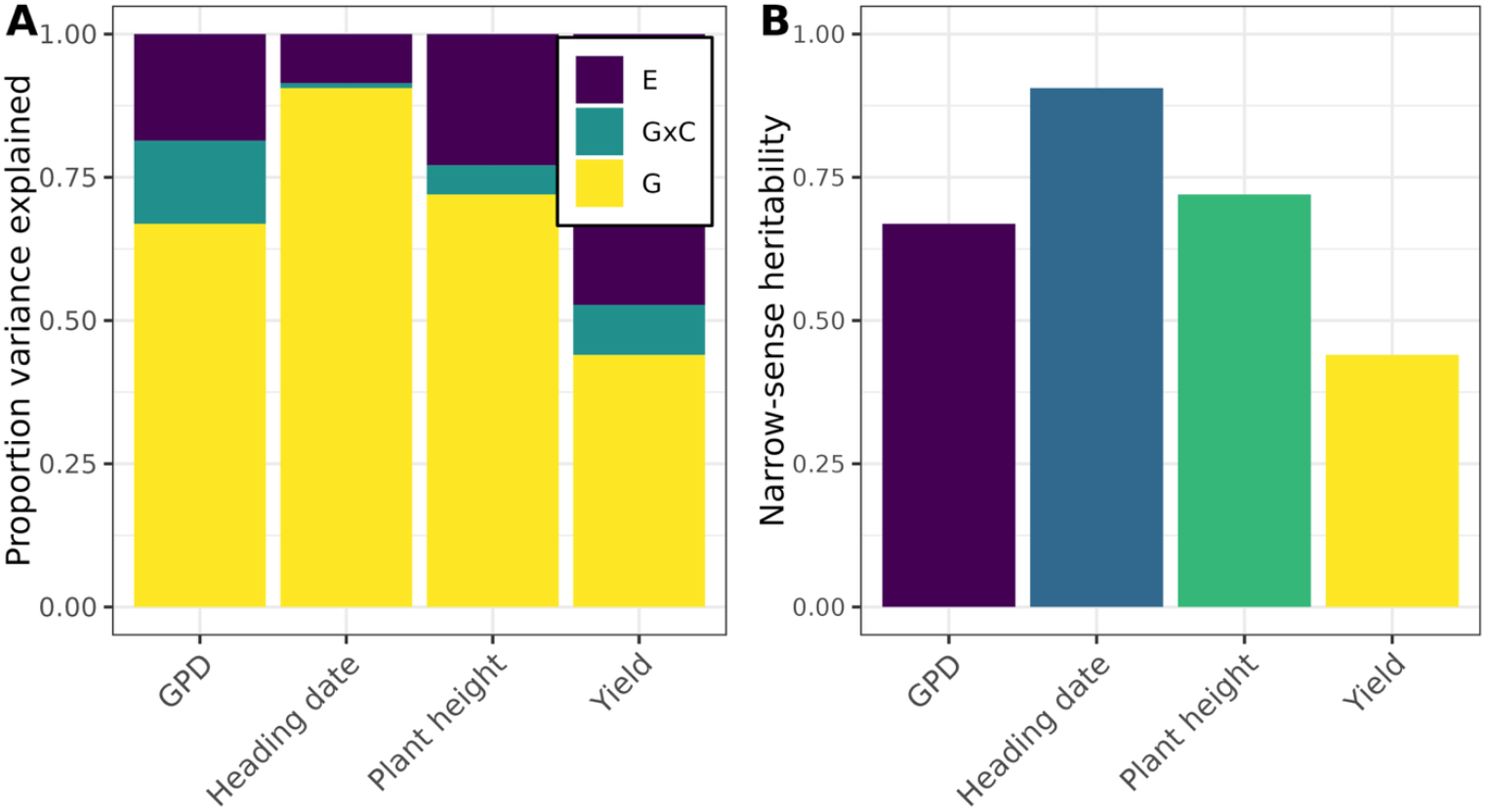
Variance components for the M1 model, and heritability estimates. A) Relative proportion of phenotypic variance explained by each variance component for each trait. Variance components from the **M1** model, where **G** is the additive genomic variance, **GxC** refers to the variance captured by the interactions between the additive genomic values and country, and **E** is the error variance. B) Narrow-sense heritability h^2^ shown for each trait, as estimated with the M1 model.

**Figure S3:**
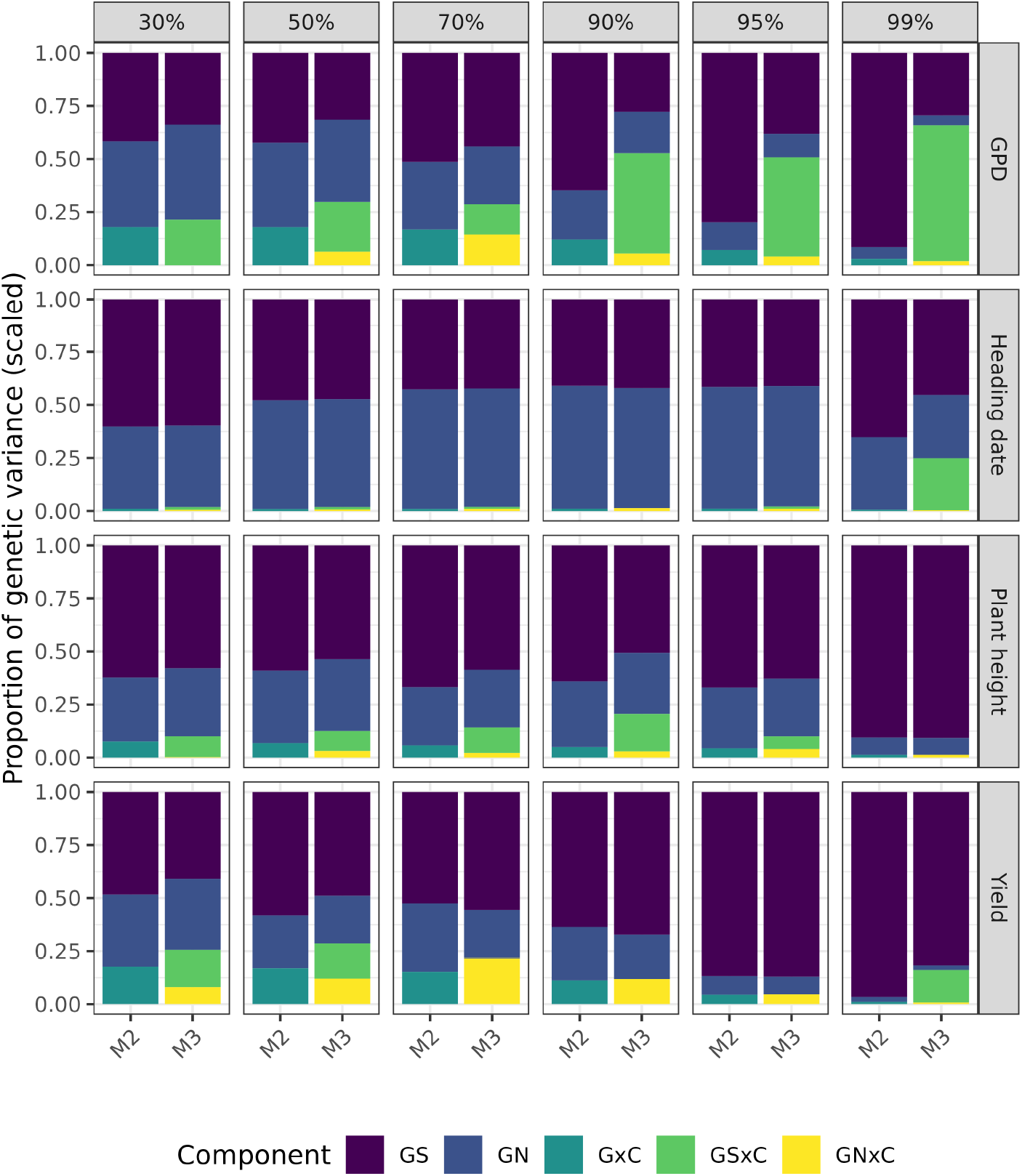
Variance components for the selection informed random effect models. Relative proportion of genetic variance explained by each of the variance components for each trait at different Z-score thresholds *z* (minimum lower percentile thresholds on absolute values) used to partition variants into selected and nonselected classes based on the inferred signatures of selection. Variance components are shown for the M2 (marginal effects of prioritized variants) or M3 models (environment-specific effects of prioritized variants). **G**_**S**_ and **G**_**N**_ represent the additive genetic effects of selected and non-selected variants, respectively. **G**_**S**_**xC** and **G**_**N**_**xC** represent their respective genotype-by-environment (country) interaction effects, while **GxC** refers to the interactions of additive genetic effects (unpartitioned) with environment (country).

